# PhaGAMeToo: A semi-automated workflow for merging structural and functional annotation of phage genomes and generation of a GenBank file

**DOI:** 10.64898/2026.08.09.738482

**Authors:** Ebrar Demircioğlu, Martin Bole, Ulisses Nunes da Rocha, René Kallies

## Abstract

**Motivation:** Analysing and concatenating phage annotation is time-consuming. Further, the output of phage annotation tools cannot be directly submitted to public repositories. To deal with these issues, we developed PhaGAMeToo. This command-line workflow for Linux integrates the functional annotations of two major viral annotation tools (Pharokka and VIBRANT), enabling faster and more accurate functional annotation. Furthermore, the workflow provides merged annotations as submission-ready GenBank files.

**Results:** PhaGAMeToo uses three steps to generate submission-ready GenBank files. The user uses the reoriented viral genomes as inputs for Pharokka and VIBRANT. Pharokka and VIBRANT-generated files are parsed through the PhaGAMeToo workflow to produce a merged GenBank file. Further, PhaGAMeToo also enables the use of BLASTP to annotate hypothetical proteins not identified by Pharokka and VIBRANT. It then merges the results into a submission-ready GenBank file(s). We tested PhaGAMeToo in three different Use Cases. We analysed reference and uncultivated viral genomes manually curated or directly recovered using MuDoGeR in our Use Cases. In the Use Case 1, we analysed four different NCBI reference genomes. In the Use Cases 2 and 3, we analysed seven recently described huge phage genomes and 56 uncultivated viral genomes recovered from 30 soil metagenomes, respectively.

**Availability and implementation:** The source code, documentation, and installation instructions for PhaGAMeToo are available at https://github.com/NFDI4Microbiota/PhaGAMeToo

**Supplementary information:** Supplementary data will be made available upon publication.

## 1 Introduction

Bacteriophages are relevant for ecology, as they are key players in the functioning of every ecosystem. At the same time, their functional capacity has great potential for biotechnology and medicine (Haq *et al*., 2012). Therefore, understanding their functions better may open new doors for ecology and biotechnology. With the advancement of viral genome recovery from metagenomes (DNA phages) (Nayfach et al., 2021) and metatranscriptomes (RNA phages) (Starr et al., 2019), the identification of novel phages is growing exponentially. However, the annotation of these genomes remains a challenge. Many phage genes remain hypothetical, as homologous reference sequences are often lacking (Wan *et al*., 2021). Beyond the presence of hypothetical proteins, phage genome annotation remains a challenge as they may have large insertions and deletions leading to gene rearrangements (Casjens, 2003), and they may have multiple gene variants or gene families (Brüssow and Hendrix, 2002).

Several workflows and software have recently been developed to make phage genome annotation easier. These existing tools are based on different open-reading frame (ORF) calling solutions. This issue can lead to different annotations for identical coding sequences (CDS) when more than one database or algorithm is used. However, a combination of tools is recommended for the best phage genome annotation (Wu et al., 2024). Such a combination requires data comparison and merging. There is a growing number of projects involving the identification of novel phages in the omics era, which need to be annotated and deposited in public repositories. If merging and comparison are done manually, the process is time-consuming, error-prone, and unsuitable for the parallel annotation of genomes in the era of Big Data. Additionally, several CDS remain hypothetical even after using different viral gene annotation tools, adding to the results’ complexity and ambiguity (Sivashankari and Shanmughavel, 2006). This issue still needs to be fully addressed in the field of virology. Furthermore, various consortia and funding agencies advocate for the FAIR deposition of data, presenting a challenge to generate a machine- and human-readable format of data files. These and their corresponding metadata must be deposited in public repositories (e.g., NCBI, ENA, DDBJ), which is an arduous process when dealing with several thousands of viral genomes. Current tools do not provide submission-ready, machine- and human-readable files prepared for data submission at the end of viral genome analysis and annotation. Our team used VIBRANT (Kieft *et al*., 2020) and Pharokka (Bouras *et al*., 2023) to annotate huge phage genomes comprehensively (Kallies et al., 2023) and encountered some of the problems mentioned above.

To deal with these issues, we designed a semi-automatic workflow that compares and combines the output of different viral annotation tools. Our workflow, the Phage Genome Annotation Merge Tool (PhaGAMeToo), is based on two tools developed for analysing and annotating phage genomes. VIBRANT is an all-round software tool for phage genome analysis. In addition to genome recovery from metagenomes and quality analysis, this software can annotate phage genomes. It uses Hidden Markov Profiles from the Kegg (Kanehisa *et al*., 2016), Pfam (Finn *et al*., 2014) and VOG (Grazziotin *et al*., 2017) databases. The second tool, Pharokka, uses the PHROGs database (Terzian *et al*., 2021) for functional annotation and also provides the ability to identify tRNAs, tmRNAs, CRISPR sequences, virulence factors and antibiotic resistance genes. The tool merges viral functional annotation results of VIBRANT and Pharokka, and additionally attempts to annotate the hypothetical CDS with a BLASTP analysis (Altschul et al., 1997). The BLASTP analysis is conducted against user-defined database(s). The final output of PhaGAMeToo is a text file, CSV files and a submission-ready GenBank file with the combined annotation results. Additionally, the tool also generates three separate folders containing the BLASTP results. PhaGAMeToo can be used to annotate single genomes or multiple genomes simultaneously. We tested PhaGAMeToo using three Use Cases. The first Use Case consisted of four reference phage genomes of different sizes collected from NCBI. For the second Use Case, we tested the pipeline using seven recently described huge phage genomes (Kallies et al., 2023). The final Use Case consisted of 56 uncultivated viral genomes (uVIGs) recovered from 31 soil metagenomes using MuDoGeR (da Rocha and Kasmanas et al., 2023).

## 2 Implementation

Users wishing to merge the functional annotations of viral genomes should use PhaGAMeToo only after annotating their respective sequences, first using the Pharokka tool (Bouras et al., 2023) and then the VIBRANT tool (Kieft et al., 2020), in this exact order. Before proceeding, the user must screen the viral sequences for the large terminase subunit (*terL*), which can be done with Pharokka. The user can either manually identify the large terminase subunit from the general feature format file (.gff) or GenBank file (.gbk) after running Pharokka. If a *terL* large terminase subunit has been identified, the user should use the terminase large subunit reorientation mode of Pharokka to reorient the genome. This mode reorients the phage to start with the large terminase subunit and then performs the annotation. Alternatively, Pharokka offers a -*-dnaapler* flag that calls the tool Dnaapler (Bouras et al., 2024), which uses BLASTX alignment against an amino acid sequence database of *terL* genes curated using PHROGs (Terzian et al., 2021) to find the desired starting sequence. The -*-dnaapler* flag reorients the genome and performs the annotation in a single step, without additional user input. At this point, the user will have a FASTA (.ffn) file and an annotated GenBank (.gbk) file, both reoriented to start with the terL gene as the first CDS, generated by Pharokka. Next, the user should use the reoriented FASTA files output by Pharokka as input files for the VIBRANT tool. This process will yield a second set of files generated by VIBRANT: a FASTA (.ffn) file and an annotated GenBank (.gbk) file. Once these files have been generated, the user needs to install the BLAST+ tool and download the required databases. After BLAST+ is appropriately installed, the user should ensure that the files are standardised so that they are compatible with the PhaGAMeToo workflow; that is, the headers of the FASTA files must match the “locus tag” regions of the corresponding GenBank files. The workflow also requires a specific file structure to be established. For each sample, the *pharokka.gbk and pharokka.ffn* output files from Pharokka need to be placed in subdirectories within the parent *pharokka/* directory, with each subdirectory named after the sample. The .*gbk and .ffn* output files from VIBRANT need to be placed directly in the *VIBRANT/* parent directory for all samples, with each file prefixed by the sample name and the appropriate extension. Once these prerequisites are met, users may run the PhaGAMeToo workflow.

We designed PhaGAMEToo to run as a Linux command line workflow with three custom scripts written in Python 3.10. The main script, *PhaGAMeToo.py*, runs the whole workflow, calling the *progress.py* for the additional BLASTP annotation, while *locustag_fix.py* standardises and formats the fasta and GenBank files and ensures compatibility with the workflow. The *progress.py* script allows for multi-core parallelisation of the BLAST process, facilitating the functional annotation of large numbers of genomes simultaneously. The user can set the number of cores used for additional annotation with a flag. System requirements for the workflow depended on the number and size of viral genomes to be processed and the databases used for BLAST alignments.

Besides the prerequisite steps (described above), PhaGAMeToo workflow consists of three main steps. The first step checks the Pharokka and Vibrant input .gbk and .ffn files and replaces any blank or non-ASCII characters with underscores (*locustag_fix*.py script). The workflow (*PhaGAMeToo.py* script) then identifies the CDS of the genome annotations from Pharokka and Vibrant and uses the locus tag and nucleotide sequence information, checking the number, length, and orientation of the CDS for inconsistencies. CDS that are predicted by only one annotation tool are included; as such, if no CDS is predicted in the same genomic region by the other tool. For CDS matching at the same genomic location but differing in length, the longest open reading frame is selected as the final product.

In the second step (*PhaGAMeToo.py* script), the workflow compares the functional assignments of each CDS. If a CDS is assigned a function from both annotation tools, it is included in the merged result. In the case of ambiguous functional assignments (e.g., major capsid protein versus major head protein), both functions are kept and separated by a slash in the output text file to allow the user to make an exact assignment individually if necessary. If only one of the tools assigns a function to a CDS, that function is transferred to the output file. If neither of the tools can assign a function, the CDS is labelled as “hypothetical protein” in the output file. PhaGAMeToo creates preliminary merged GenBank files based on the final functional annotations presented in the CSV files.

All hypothetical proteins are automatically subjected to a BLAST analysis using BLASTP alignments in the third step (*progress.py* script) against a user-provided database. Different databases can be used based on available local space. The Swissprot and Refseq-protein databases are most suited for local usage (i.e., laptops) of the BLAST tool, as they take up <1 Gb and 5 Gb of space, respectively. Using BLAST with the NCBI non-redundant database is more suitable for High-Performance Cluster (HPC) or workstation analysis, as its size, at the time of writing, is 563 GB and likely to increase in size over time.

All BLASTP results are provided separately for each database used in a folder but are not included in the output GenBank file. For example, when blasting against the NCBI non-redundant database, the first hits might appear as hypothetical proteins, as the next hits are against poorly annotated phages, with only a hit with less similarity leading to a function. The automatic transfer of such results into the final output file leads to incorrect results. Therefore, users must manually evaluate the BLASTP results and, if necessary, transfer them to the final output file.

We encourage our end users to use multiple databases for the functional annotation of genomic sequences to ensure accurate and comprehensive results. To date, no single database encompasses all currently known functional annotations, and specialised tools for viral annotation are constantly being developed. Thus, it is safe to assume that using multiple databases for annotation enhances the reliability and robustness of functional predictions while providing a way to cross-validate the results.

## 3 Application

We validated our workflow using three datasets. The first, Use Case 1, consisted of four reference phage genomes with different genome sizes, i.e. *Escherichia* phage phiX174 (5,386 bp ss-DNA, GenBank Acc. no. NC_001422.1), *Escherichia* phage MS2 (3,569 bp ss-RNA, Genbank Acc. No. NC_001417.2), *Enterobacteria* phage T4 (168,903 bp ds-DNA, GenBank Acc. No. NC_000866.4) and Prevotella phage Lak-C1 (540,217 bp ds-DNA, GenBank Acc.No. MK250029.1). All genome annotations were merged with PhaGAMEToo, and compared with results done using manual annotation. Our analyses showed that, for Use Case 1, we identified 1,75% more CDS annotation using PhaGAMeToo automated workflow than when we analysed the different viral genomes manually (Supplementary Table S1, Supplementary Files S1 - S4).

For Use Case 1, encompassing the four reference phage genomes, the entire workflow was executed in a few seconds on a 6-core, 11^th^-generation Intel Core i5-11400H processor computer running Kali GNU/Linux 2023.4 Rolling Release. For further analysing the generation of GenBank files manually and using PhaGAMEToo, we designed the Use Case 2.

The scientific results for Use Case 2 and Use Case 3, have been computed at the High-Performance Computing (HPC) Cluster EVE, a joint effort of both the Helmholtz Centre for Environmental Research - UFZ (http://www.ufz.de/) and the German Centre for Integrative Biodiversity Research (iDiv) Halle-Jena-Leipzig (http://www.idiv-biodiversity.de/). The workflow was executed on 56 CPU cores (dual-socket Intel Xeon 6348 CPUs), allocating 8 GB of RAM (DDR4) per CPU core for each execution, respectively.

To make sure we would have a fair comparison between manually curated and PhaGAMEToo GenBank files, Use Case 2, we used seven huge phage genomes recovered from wastewater metagenomes, where the functional annotation of said genomes was done manually (Supplementary Files S5 - S11) (Kallies *et al*., 2023). These seven genomes were used as batch input for PhaGAMeToo. When comparing the huge phage genomes annotated with PhaGAMeToo, we found 2993 functional CDS (2815), tRNA (158) and pseudogene (20) sequences, as opposed to 3028 CDS (2859) and tRNA (169) sequences as described previously (Kallies et al., 2023) and (Supplementary table S2, Supplementary file S20 - S21, Supplementary Figure S1-S3). This 1,16 % difference is due to presenting only the raw merged annotations from the Pharokka and VIBRANT tools, without manual curation and inclusion of BLAST annotated hypothetical proteins in the results (Supplementary table S2, Supplementary file S20 - S21, Supplementary Figure S1-S3). Compared to the manual analysis of the genomes, which took several hours per genome, the analysis with PhaGAMeToo was completed within a few minutes. To determine the merging time, we ran the workflow 10 times, using the same HPC node each time. On average, the merging procedure took 87.05 seconds (Supplementary Table S3). In addition, the automated nature of PhaGAMeToo significantly reduced user involvement, streamlining the workflow and reducing the risk of error when researchers manually transfer functional annotations between files.

We designed Use Case 3 to evaluate PhaGAMEToo in a novel metagenome dataset and to test PhaGAMEToo in a HPC. This dataset consisted of 56 complete or high-quality uVIGs (Supplementary Table S4) recovered from 31 soil metagenomes using MuDoGeR (da Rocha and Kasmanas et al., 2023). We utilised the NCBI non-redundant, SwissProt, and RefSeq-protein databases for alignment, performing analyses with each database individually and all three combined simultaneously. We tested the workflow on the entire dataset (of 56 uVIGs) or smaller batches. Batches were prepared in sets of three, for five, ten, twenty, thirty, forty and fifty samples, respectively, utilising the *random* module from the random Python library, setting the *seed(123)*. The 56 uVIGs contained 2782 hypothetical proteins that required annotation (73.9% out of the 3776 CDS regions), while the batches had a range of hypothetical proteins spanning from 214 (for five samples) to 2502 (for 50 samples) (Supplementary file 6). Metrics like CPU time per run, CPU utilisation, max RAM consumed and elapsed time were collected for each execution (Supplementary Table 13). We estimated how long the merging and annotation process takes for tested databases using these metrics. We created a linear regression model with interactions using robust standardised errors (Supplementary Files S12 - S19, Supplementary Files S22 - S25, Supplementary Figure S4). The model indicated that using the configuration mentioned above, it takes 1969 seconds (approximately 33 min) per hypothetical protein to annotate it using the NCBI non-redundant database, 361 seconds (approximately 6 min) for the Refseq-protein database, 1.6 seconds to annotate with Swissprot database and 2333 seconds (approximately 39 min) when using all three of the databases above Supplementary Files S12 - S19, Supplementary Files S22 - S25, Supplementary Figure S4). Using the Max RAM consumed during the annotation process, we estimated the RAM requirements for the workflow execution and rounded it up to the next whole integer. We concluded that for annotation using Swissprot and Refseq-protein, 1 GB of RAM per CPU core is sufficient, while for the NCBI non-redundant database and all three databases simultaneously, 5 GB of RAM per CPU core is required (Supplementary File S15 - S16).

We recommend that, in case the user would like to merge and create the submission-ready GenBank files and additional BLAST annotation files for several hundreds or thousands of viral or uVIG samples, they split their datasets into batches such that there are 3000 - 3500 hypothetical proteins per batch.

Compared to the manual functional annotation merging process, which is often reported to take several days to weeks, PhaGAMeToo offers significant time advantages while eliminating the possibility of erroneous merging. Therefore, PhaGAMETOO increases the accuracy and speed of phage annotation, making our workflow ideal for large-scale genetic research, where speed, precision, and consistency are essential.

## 4 Conclusion

PhaGAMeToo is a semi-automated workflow for efficient phage genome annotation. It seamlessly merges the findings of two standard phage genome annotation tools, Pharokka and VIBRANT. PhaGAMeToo uses these two tools but could be adapted to other tools to optimise the phage genome annotation process. By merging data from Pharokka and VIBRANT, PhaGAMeToo improves the functional annotation process speed, completeness and accuracy. Users benefit from PhaGAMeToo’s ability to effortlessly generate final functional annotations for each phage genome and generate new GenBank files based on these annotations, including the possibility to enhance the annotation of hypothetical proteins further. The simplicity of a single-command operation makes PhaGAMeToo highly accessible and facilitates its use by scientists with limited bioinformatics expertise.

## Supporting information

Supplementary Information (legends for supplementary files)

## Acknowledgement

Assoc. Prof. Dr. Kasım Öztoprak, Department of Computer Engineering, Konya Food and Agriculture University, for providing high-performance computing capacities.

## Funding

This work was funded by the Deutsche Forschungsgemeinschaft under NFDI4Microbiota consortium, grant number 460129525, and the Helmholtz Young Investigator, grant VH-NH-1248 Micro’ BigData’.

---

**Figure 1:**
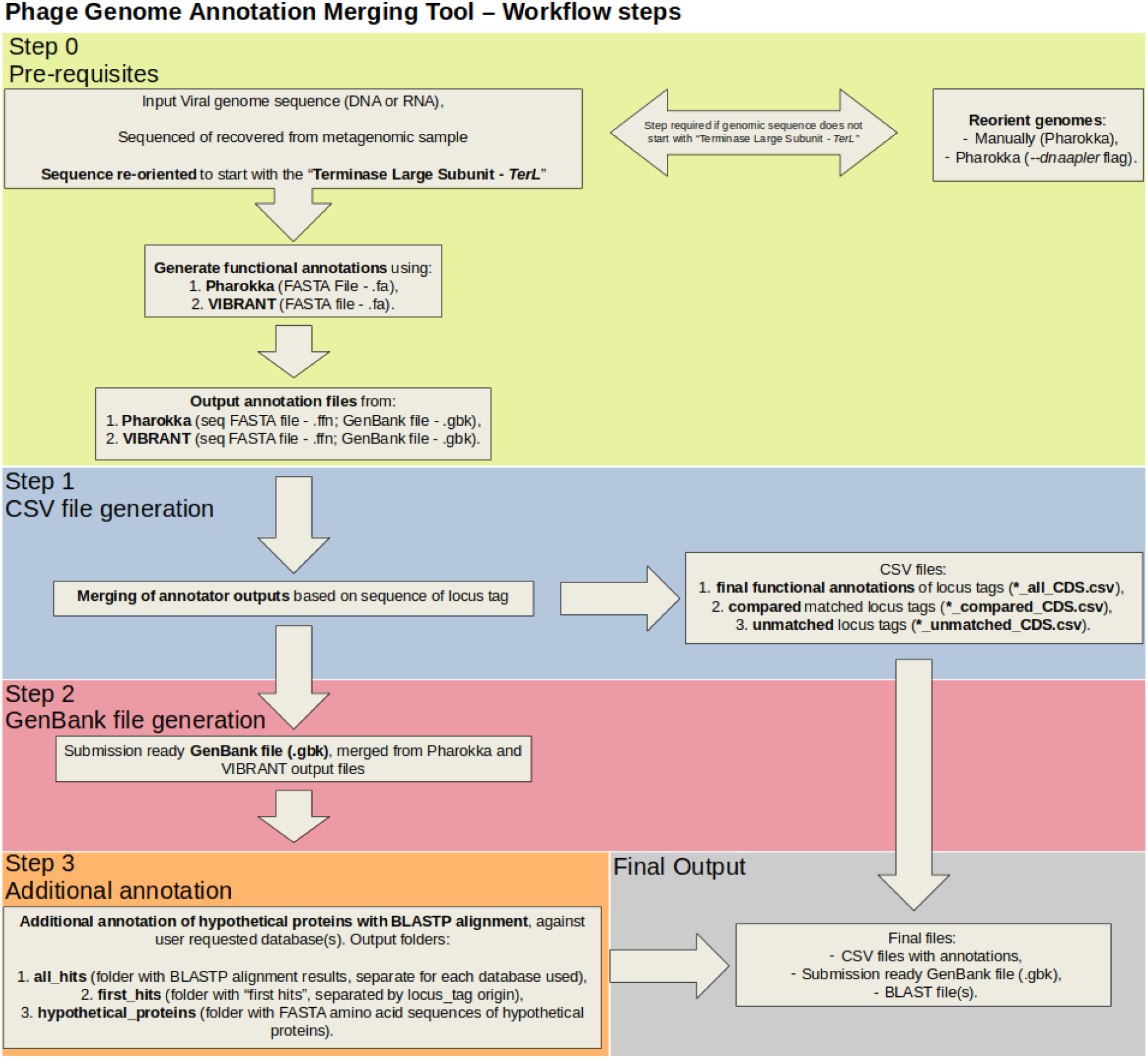
PhaGAMeToo workflow. The tool starts with the Pharokka and Vibrant input gbk files and checks the number, length and orientation of the CDS generated by each tool. The second step checks and merges the functional assignments of each CDS in the phage genome and generates output files. Additional automated BLAST analyses of hypothetical protein sequences are performed in a third step, allowing users to improve the annotation of the phage genome.

## Notes

https://github.com/NFDI4Microbiota/PhaGAMeToo

### Competing Interest Statement

The authors have declared no competing interest.

