## Supplementary Information (legends for supplementary files) for "PhaGAMeToo: A semi-automated workflow for merging structural and functional annotation of phage genomes and generation of a GenBank file"

Manuscript ID BIORXIV/2026/738482

### Scope

This document is the legend for the supplementary files accompanying the manuscript. The files provide the measurements and comparative analyses on which the manuscript's statements about annotation completeness, runtime and memory requirements are based. They are presented in two groups: accuracy against manual annotation, and performance and resource requirements.

Identifiers and filenames are those of the supplementary file set referenced in the manuscript text. Nineteen files are included, totalling approximately 4.6 MB.

Throughout, a hypothetical protein is a coding sequence for which the annotation tool returned no functional assignment. Kallies et al. (2023) refers to the manually curated annotation of seven wastewater phage genomes published in Viruses 15:2330, which serves as the manual reference standard in Group 1.

### Group 1. Accuracy against manual annotation (Use Case 2)

Seven high-molecular-weight phage genomes recovered from wastewater metagenomes were annotated independently by Pharokka, by VIBRANT, by manual curation (Kallies et al., 2023) and by PhaGAMeToo. The files in this group report that four-way comparison.

#### Supplementary Table S2

**Filename:** Supplementary_table_S2_20241021_tool_comparison_table.csv

**Format:** Comma-separated values, UTF-8

**Size:** 1.3 kB (8 data rows, 17 columns)

Per-genome comparison of annotation completeness across the four approaches. Rows are the seven genomes, Genbank_1-SewaA to Genbank_7-SewaG, followed by a total row summing each numeric column; percentages in the total row are recomputed from the totals rather than averaged across rows.

| **Column** | **Description** |
| --- | --- |
| sample name | Genome identifier, Genbank_{n}-Sewa{X}, or 'total' for the summary row. |
| Number of CDS after Kallies et al | Coding sequences annotated by manual curation. |
| Number of tRNA after Kallies et al | Transfer RNA genes annotated by manual curation. |
| Number of pseudogene after Kallies et al | Pseudogenes annotated by manual curation; zero throughout, as the manual procedure did not assign this category. |
| Number of hypo after Kallies et al | Coding sequences left without functional assignment after manual curation. |
| Percentage of hypothetical proteins after Kallies et al | The preceding column as a percentage of manually annotated coding sequences. |
| Number of CDS PhaGAMeToo | Coding sequences in the PhaGAMeToo merged annotation. |
| Number of tRNA PhaGAMeToo | Transfer RNA genes in the PhaGAMeToo merged annotation. |
| Number of pseudogene PhaGAMeToo | Pseudogenes in the PhaGAMeToo merged annotation. |
| Number of hypo protein after PhaGAMeToo | Coding sequences without functional assignment after merging. |
| Percentage of hypothetical proteins after PhaGAMeToo | The preceding column as a percentage of PhaGAMeToo coding sequences. |
| Number of locus_tag found Pharokka | Distinct locus tags in the Pharokka annotation, that is, all features called by Pharokka. |
| Number of hypo protein after Pharokka | Coding sequences without functional assignment in the Pharokka annotation. |
| Percentage of hypothetical proteins after pharokka | The preceding column as a percentage of Pharokka locus tags. |
| Number of locus_tag found VIBRANT | Distinct locus tags in the VIBRANT annotation. |
| Number of hypo protein after VIBRANT | Coding sequences without functional assignment in the VIBRANT annotation. |
| Percentage of hypothetical proteins after VIBRANT | The preceding column as a percentage of VIBRANT locus tags. |

The total row gives 2,859 CDS, 169 tRNA and 0 pseudogenes, that is 3,028 features, for the manual annotation, against 2,815 CDS, 158 tRNA and 20 pseudogenes, that is 2,993 features, for PhaGAMeToo: a difference of 1.16 per cent. Hypothetical proteins account for 78.36 per cent of manually annotated features and 76.44 per cent of PhaGAMeToo features, against 79.29 per cent for Pharokka and 79.95 per cent for VIBRANT alone.

#### Supplementary File S21

**Filename:** Supplementary_file_S21_20241021_comparative_analysis_explanation.txt

**Format:** Plain text, UTF-8

**Size:** 2.0 kB

Statistical interpretation of Supplementary Table S2. Paired t-tests across the seven genomes show that both manual curation and PhaGAMeToo reduce the number of hypothetical proteins relative to Pharokka (p = 0.02173) and relative to VIBRANT (p = 0.002759). The paired comparison between manual curation and PhaGAMeToo yields a mean difference of zero, indicating that the semi-automated merge recovers as many functional assignments as expert manual merging of the same two annotation sources.

#### Supplementary Figure S1

**Filename:** Supplementary_figure_S1_20241128_combined_comparative_analysis_plot.png

**Format:** PNG, 4200 × 2100 pixels, 8-bit RGB

**Size:** 407 kB

Two-panel bar chart. The left panel gives, for each of the seven genomes, the absolute number of coding sequences and of hypothetical proteins under each of the four annotation approaches. The right panel gives the corresponding percentage of hypothetical proteins relative to total coding sequences. Both merging approaches reduce the hypothetical fraction relative to either source tool alone, and do so by an equal amount.

#### Supplementary Figure S2

**Filename:** Supplementary_figure_S2_20241128_plot_counts_analysis.png

**Format:** PNG

**Size:** 282 kB

The absolute-count panel of Supplementary Figure S1 as a standalone figure, at a resolution permitting inspection of individual genomes.

#### Supplementary Figure S3

**Filename:** Supplementary_figure_S3_20241128_plot_perc_analysis.png

**Format:** PNG

**Size:** 209 kB

The percentage panel of Supplementary Figure S1 as a standalone figure. Percentages normalise for the large differences in genome size across the seven samples.

#### Supplementary File S20

**Filename:** Supplementary_file_S20_20241021_combined_comparative_analysis_plot_explanation.txt

**Format:** Plain text, UTF-8

**Size:** 2.0 kB

Description of the construction and interpretation of Supplementary Figures S1 to S3, including the definition of each panel and of the four annotation approaches compared.

#### Supplementary Files S5 to S11

**Filenames:** Supplementary_file_S5_Genbank_1-SewaA.txt; Supplementary_file_S6_Genbank_2-SewaB.txt; Supplementary_file_S7_Genbank_3-SewaC.txt; Supplementary_file_S8_Genbank_4-SewaD.txt; Supplementary_file_S9_Genbank_5-SewaE.txt; Supplementary_file_S10_Genbank_6-SewaF.txt; Supplementary_file_S11_Genbank_7-SewaG.txt

**Format:** GenBank flat file, plain text

**Sizes:** 630 kB, 616 kB, 474 kB, 460 kB, 694 kB, 473 kB and 521 kB respectively; 3.9 MB in total

The manually curated annotations of the seven wastewater phage genomes published by Kallies et al. (2023). These files constitute the reference standard against which the PhaGAMeToo output is compared in Supplementary Table S2 and Supplementary Figures S1 to S3, and are included so that the comparison can be verified independently.

Each file follows the standard GenBank structure: a LOCUS line giving the organism designation, sequence length, molecule type and topology; a FEATURES table in which each CDS carries locus_tag, product and translation qualifiers and each tRNA carries locus_tag and product qualifiers; and the nucleotide sequence in ORIGIN format.

### Group 2. Performance and resource requirements (Use Cases 2 and 3)

Runtime and memory consumption were measured with /usr/bin/time -v on the EVE cluster of the Helmholtz Centre for Environmental Research and the German Centre for Integrative Biodiversity Research, using nodes with 56 CPU cores (dual-socket Intel Xeon 6348) and 8 GB DDR4 RAM per core.

#### Supplementary Table S3

**Filename:** Supplementary_table_S3_20241128_merging_time_comparison_table.csv

**Format:** Comma-separated values, UTF-8

**Size:** 650 B (10 data rows, 5 columns)

Elapsed time of ten repeated executions of the merging step on the seven wastewater phage genomes of Use Case 2, all run on the same cluster node so that hardware variance does not contribute to the spread.

| **Column** | **Description** |
| --- | --- |
| Number of run | Sequential index of the replicate, 1 to 10. |
| SLURM job record | Name of the /usr/bin/time -v output file for that replicate, time_output_nr00.txt to time_output_nr09.txt. |
| JobID | Scheduler job identifier, in the form 12_WW_phages-<SLURM job number>. |
| Elapsed time | Wall-clock duration of the merging step, in seconds. |
| EVE Node | Compute node on which the replicate was executed; node024 for all ten runs. |

Elapsed times range from 84.38 to 91.31 seconds, with a mean of 87.05 seconds.

#### Supplementary File S13

**Filename:** Supplementary_file_S13_20240923_per-set_benchmark_results.csv

**Format:** Comma-separated values, UTF-8

**Size:** 8.3 kB (72 data rows, 17 columns)

One row per benchmark execution. The 72 rows are the eighteen input sets, namely three independent replicates at each of the batch sizes 5, 10, 20, 30, 40 and 50 uncultivated viral genomes, each processed against four database configurations: SwissProt, RefSeq-protein, the NCBI non-redundant database, and all three simultaneously. Column definitions are also supplied in machine-readable form as Supplementary File S14.

| **Column** | **Description** |
| --- | --- |
| set_name | Identifier of the input batch, set_{replicate}_{batch size}. |
| sample_count | Number of uncultivated viral genomes in the batch. |
| db_name | Database configuration used: swissprot, refseq-protein, nr, or allDB for all three combined. |
| CPU Time | Total user-level CPU time consumed, HH:MM:SS. |
| CPU System Time | Total kernel-level CPU time consumed, HH:MM:SS. |
| CPUs Used | Number of CPU cores effectively used, derived from the reported CPU utilisation percentage. |
| Elapsed Time | Wall-clock duration of the execution, HH:MM:SS. |
| Max RAM (GB) | Peak resident memory across the execution, in gigabytes. |
| RAM Usage Percentage | Peak resident memory as a percentage of the memory available on the node. |
| Minimal RAM Requirement | Minimum memory required per CPU core, in gigabytes, rounded up to the next integer. |
| number_of_hypothetical_proteins_processed | Number of hypothetical proteins submitted to BLASTP in that execution. This column holds the per-batch counts quoted in the manuscript, ranging from 214 for a five-genome batch to 2,502 for a fifty-genome batch. |
| CPU Time/hypothetical_protein (seconds) | User CPU time divided by the number of hypothetical proteins processed, in seconds. |
| CPU Time/hypothetical_protein (minutes) | The same quantity expressed in minutes. |
| Major Page Faults | Page faults requiring a disk read, an indicator of memory pressure. |
| Minor Page Faults | Page faults served from memory without disk access. |
| Voluntary Context Switches | Occasions on which the process yielded the CPU, typically while awaiting input or output. |
| Involuntary Context Switches | Occasions on which the kernel preempted the process. |

#### Supplementary File S14

**Filename:** Supplementary_file_S14_20240927_per-set_benchmark_columns_explanation.csv

**Format:** Comma-separated values, UTF-8

**Size:** 1.8 kB (16 data rows, 2 columns)

Column dictionary for Supplementary File S13.

| **Column** | **Description** |
| --- | --- |
| Column Name | Name of a column in Supplementary File S13. |
| Description | Definition of that column, including units where applicable. |

#### Supplementary File S15

**Filename:** Supplementary_file_S15_20240923_summaryDB_benchmark_results.csv

**Format:** Comma-separated values, UTF-8

**Size:** 2.0 kB (24 data rows, 12 columns)

Supplementary File S13 aggregated over the three replicates of each batch size and database configuration, giving one row per combination of the six batch sizes and four database configurations. This table is the direct basis for the memory recommendations stated in the manuscript: 1 GB of RAM per CPU core for SwissProt and RefSeq-protein, and 5 GB per core for the NCBI non-redundant database and for all three databases combined. Column definitions are also supplied as Supplementary File S16.

| **Column** | **Description** |
| --- | --- |
| Sample Count | Number of uncultivated viral genomes in the batches averaged. |
| Database | Database configuration: swissprot, refseq-protein, nr, or allDB. |
| Average CPU Time (HH:MM:SS) | Mean user-level CPU time over the three replicates. |
| Average CPU System Time (HH:MM:SS) | Mean kernel-level CPU time over the three replicates. |
| Average Elapsed Time (HH:MM:SS) | Mean wall-clock duration over the three replicates. |
| Average Max RAM (GB) | Mean peak resident memory, in gigabytes. |
| Average CPUs Used | Mean number of CPU cores effectively used. |
| Average RAM Usage Percentage | Mean peak memory as a percentage of node memory. |
| Average Minimal RAM Requirement | Mean minimum memory per CPU core, in gigabytes. |
| Average Number of Hypothetical Proteins Processed | Mean number of hypothetical proteins submitted to BLASTP. |
| Average CPU Time/hypothetical_protein (seconds) | Mean user CPU time per hypothetical protein, in seconds. |
| Average CPU Time/hypothetical_protein (minutes) | The same quantity expressed in minutes. |

#### Supplementary File S16

**Filename:** Supplementary_file_S16_20240927_summaryDB_benchmark_results_explanation.csv

**Format:** Comma-separated values, UTF-8

**Size:** 1.5 kB (12 data rows, 2 columns)

Column dictionary for Supplementary File S15.

| **Column** | **Description** |
| --- | --- |
| Column Name | Name of a column in Supplementary File S15. |
| Description | Definition of that column, including units where applicable. |

#### Supplementary Table S4

**Filename:** Supplementary_table_S4_20241128_UViGs_IDs_provenance_taxonomy.csv

**Format:** Comma-separated values, UTF-8

**Size:** 21 kB (56 data rows, 28 columns)

The 56 complete or high-quality uncultivated viral genomes (uVIGs) recovered from 31 soil metagenomes that constitute the benchmark dataset of Use Case 3, one row per genome. The genomes were recovered with MuDoGeR v1.0.1; viral sequences were identified with VirSorter2 v2.2.4, VirFinder v1.1 and VIBRANT v1.2.1 and dereplicated with Stampede-clustergenomes; sequence quality was assessed with CheckV v1.0.1; and taxonomy was predicted with vConTACT3 v3.0.0.b65. Empty cells indicate that the corresponding value was not assigned by the recovery pipeline.

| **Column** | **Description** |
| --- | --- |
| lib_and_contig_id | Concatenated library and contig identifier, unique per genome. |
| Lib_ID | Identifier of the sequencing library from which the genome was assembled. |
| Contig_ID | Assembly contig name, encoding contig number, length and k-mer coverage. |
| contig_length | Length of the contig in base pairs. |
| genome_copies | Number of copies of the genome detected in the assembly. |
| gene_count | Total number of genes predicted on the contig. |
| viral_genes | Number of predicted genes assigned to viral hallmark or viral-like families. |
| host_genes | Number of predicted genes assigned to cellular host families. |
| checkv_quality | Genome quality tier assigned by CheckV, for example Complete or High-quality. |
| miuvig_quality | Quality tier under the Minimum Information about an Uncultivated Virus Genome standard. |
| completeness | Estimated genome completeness, in per cent. |
| completeness_method | Method used to estimate completeness, for example AAI-based. |
| contamination | Estimated proportion of non-viral sequence, in per cent. |
| prophage | Whether the sequence was identified as integrated in a host genome (Yes or No). |
| termini | Terminal repeat structure detected, for example 53-bp-DTR for a direct terminal repeat, or No. |
| GenomeName | Genome name where assigned by the taxonomy step; empty where none was assigned. |
| RefSeqID | RefSeq accession of the closest reference, where one was identified. |
| Proteins | Number of protein-coding sequences used in the taxonomic assignment. |
| Reference | Whether the genome served as a reference in the clustering step (True or False). |
| Size (Kb) | Genome length in kilobases. |
| realm (prediction) | Predicted taxonomic realm, for example Duplodnaviria, or No Realm where unassigned. |
| phylum (prediction) | Predicted phylum, for example Uroviricota. |
| class (prediction) | Predicted class, for example Caudoviricetes. |
| order (prediction) | Predicted order. Names of the form novel_order_* denote candidate lineages without an established name; vertical bars separate equally supported alternatives. |
| family (prediction) | Predicted family, following the same conventions. |
| subfamily (prediction) | Predicted subfamily, following the same conventions. |
| genus (prediction) | Predicted genus, following the same conventions. |
| network | Identifier of the gene-sharing network component to which the genome was assigned. |

### Files cited in the manuscript but not included

Supplementary Table S1 and Supplementary Files S1 to S4, cited in connection with Use Case 1, are not part of this submission. The comparison reported for those four reference genomes, namely Escherichia phage phiX174 (NC_001422.1), Escherichia phage MS2 (NC_001417.2), Enterobacteria phage T4 (NC_000866.4) and Prevotella phage Lak-C1 (MK250029.1), is stated in the manuscript text.

### Data and code availability

The source code, documentation and installation instructions for PhaGAMeToo are available at https://github.com/NFDI4Microbiota/PhaGAMeToo under the GNU General Public License version 3.
