## Supplementary Tables S2-S4, Figures S1-S3 and Files S5-S16, S20-S21 for "PhaGAMeToo: A semi-automated workflow for merging structural and functional annotation of phage genomes and generation of a GenBank file": Supplementary_file_S20_20241021_combined_comparative_analysis_plot_explanation.txt

Supplementary Figure S1: Comparative Analysis of Hypothetical Protein Annotations by Sample

This figure consists of two bar charts placed side by side, presenting a comparison of hypothetical protein annotations across different samples and methods. The left panel shows the absolute counts of coding sequences (CDS) and hypothetical proteins, while the right panel presents the percentages of hypothetical proteins relative to the total CDS for each sample.

In the left panel, the bars represent the total number of CDS and hypothetical proteins identified by four different methods: Pharokka, VIBRANT, Kallies et al. (2023; manual merging), and PhaGAMeToo (semi-automated merging). For each sample, the counts are visualized to highlight the reduction in hypothetical proteins after the merging processes. A lower count of hypothetical proteins in the Kallies et al. (2023) and PhaGAMeToo methods compared to Pharokka and VIBRANT suggests an improvement in protein annotation through the merging process.

The right panel shows the percentage of hypothetical proteins for each sample after annotation, allowing for a relative comparison of the methods' performance. The reduction in the percentage of hypothetical proteins after the merging (Kallies et al. (2023) and PhaGAMeToo) compared to the initial annotation methods (Pharokka and VIBRANT) further demonstrates the effectiveness of these approaches in improving the annotation process.

Overall, the figure indicates that both Kallies et al. (2023) and PhaGAMeToo significantly reduce the number of hypothetical proteins compared to Pharokka and VIBRANT. The fact that the two merging methods yield identical results across all samples suggests that both are equally effective at improving annotation quality by minimizing unannotated hypothetical proteins. These figures do not take into account the fact that manual curation is a long and error-prone process, while the semi-automated workflow is executed in mere seconds.
