## Supplementary Tables S2-S4, Figures S1-S3 and Files S5-S16, S20-S21 for "PhaGAMeToo: A semi-automated workflow for merging structural and functional annotation of phage genomes and generation of a GenBank file": Supplementary_file_S21_20241021_comparative_analysis_explanation.txt

Supplementary Table S2: Comparison of Hypothetical Protein Annotation 

In this analysis, we compared the effectiveness of four different ways to comprehensively annotate uncultivated viral genomes (uVIGs), or more specifically, their coding regions (CDS). We looked at the total number of CDS (including tRNA and pseudogene regions). After annotation by their respective tools or workflows, we compared it to the total number of hypothetical proteins after annotation. The goal of the comparison was to assess whether the manual merging method employed by Kallies et al. (2023) and the semi-automated merging workflow by PhaGAMeToo could significantly reduce the number of hypothetical proteins compared to the original annotations from Pharokka and VIBRANT, thus providing a more detailed insight into the functional potential of viral sequences.

Using paired t-tests, we found that both Kallies et al. (2023) and PhaGAMeToo significantly reduced the number of hypothetical proteins compared to Pharokka (p-value = 0.02173) and VIBRANT (p-value = 0.002759). The mean difference indicated that Pharokka and VIBRANT resulted in fewer functional annotations of CDS regions than both Kallies et al. (2023) and PhaGAMeToo, respectively. Furthermore, the paired t-test between Kallies et al. (2023) and PhaGAMeToo revealed no difference in the number of hypothetical proteins between the two methods (mean difference = 0), suggesting that both methods are equally effective in reducing the number of hypothetical proteins.

Overall, the results suggest that both the manual merging process as employed by Kallies et al. (2023) and the semi-automated PhaGAMeToo workflow offer significant improvements over Pharokka and VIBRANT in reducing the number of hypothetical proteins, with no statistical difference between the manual and semi-automated merging methods. This analysis does not take into account the fact that manual curation is a long and error-prone process, while the semi-automated workflow is executed in mere seconds.
