## Supplementary figures and images for "PhaGAMeToo: A semi-automated workflow for merging structural and functional annotation of phage genomes and generation of a GenBank file"

### Supplementary_figure_S1_20241128_combined_comparative_analysis_plot.png

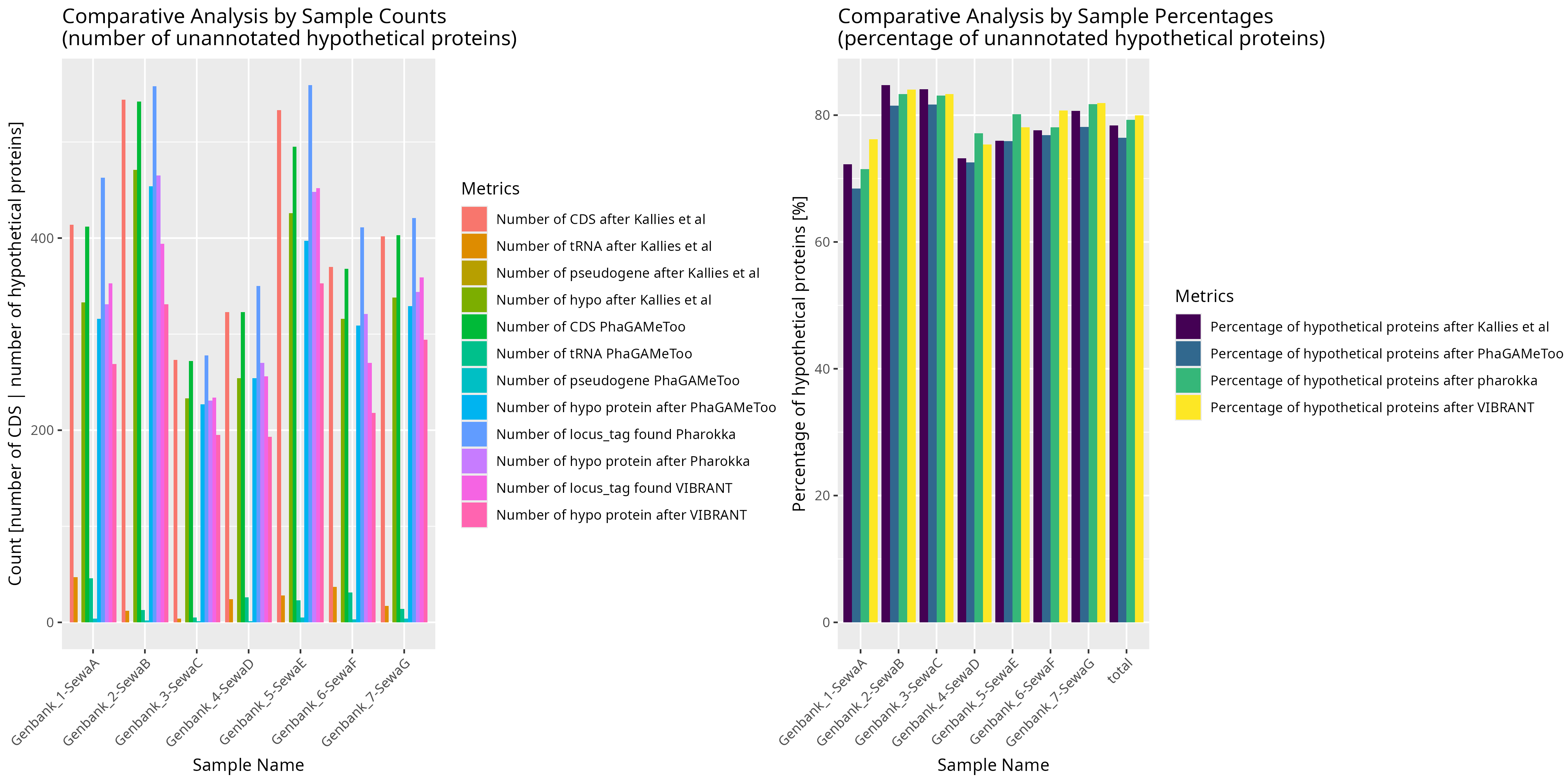

### Supplementary_figure_S2_20241128_plot_counts_analysis.png

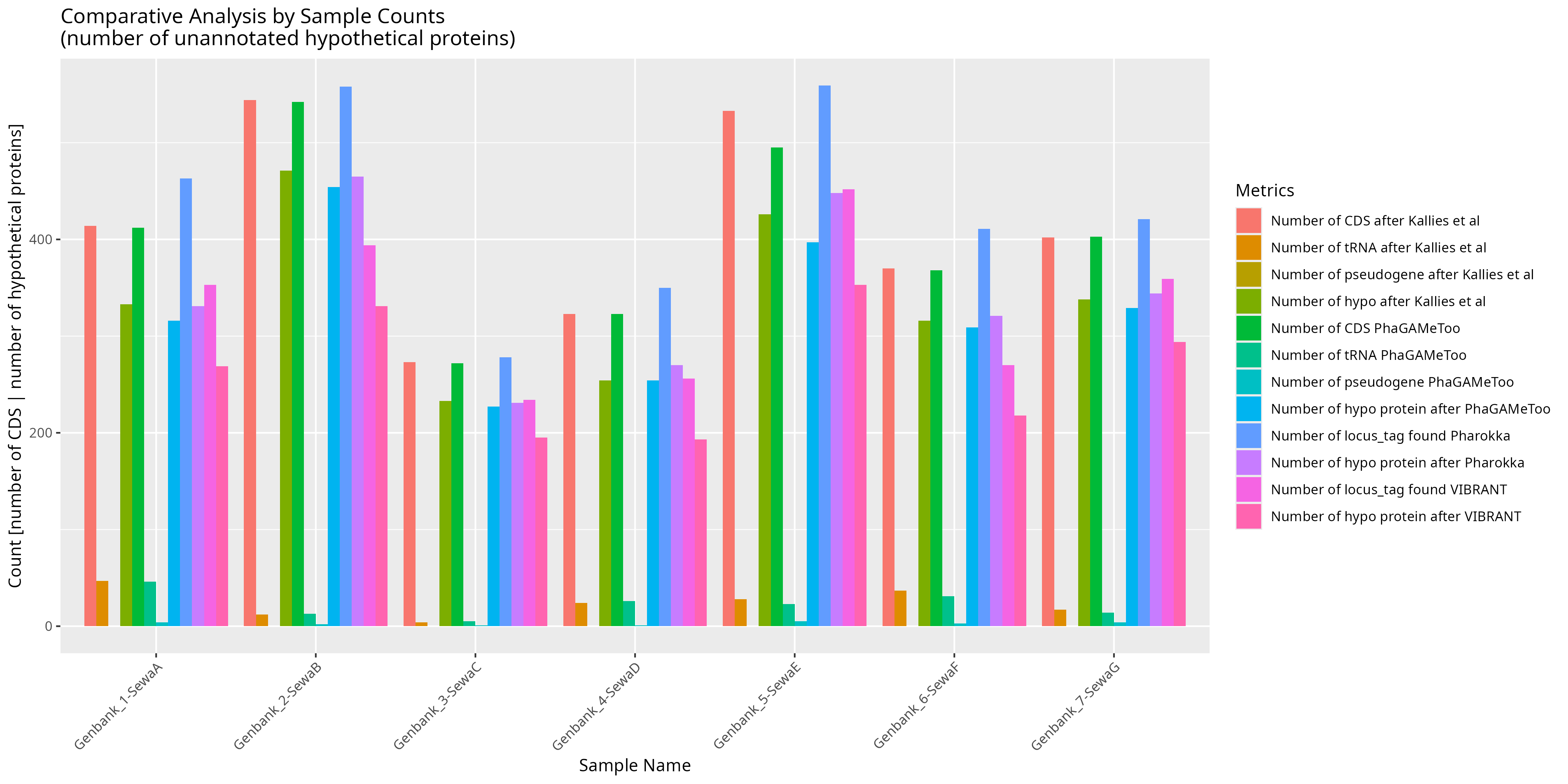

### Supplementary_figure_S3_20241128_plot_perc_analysis.png

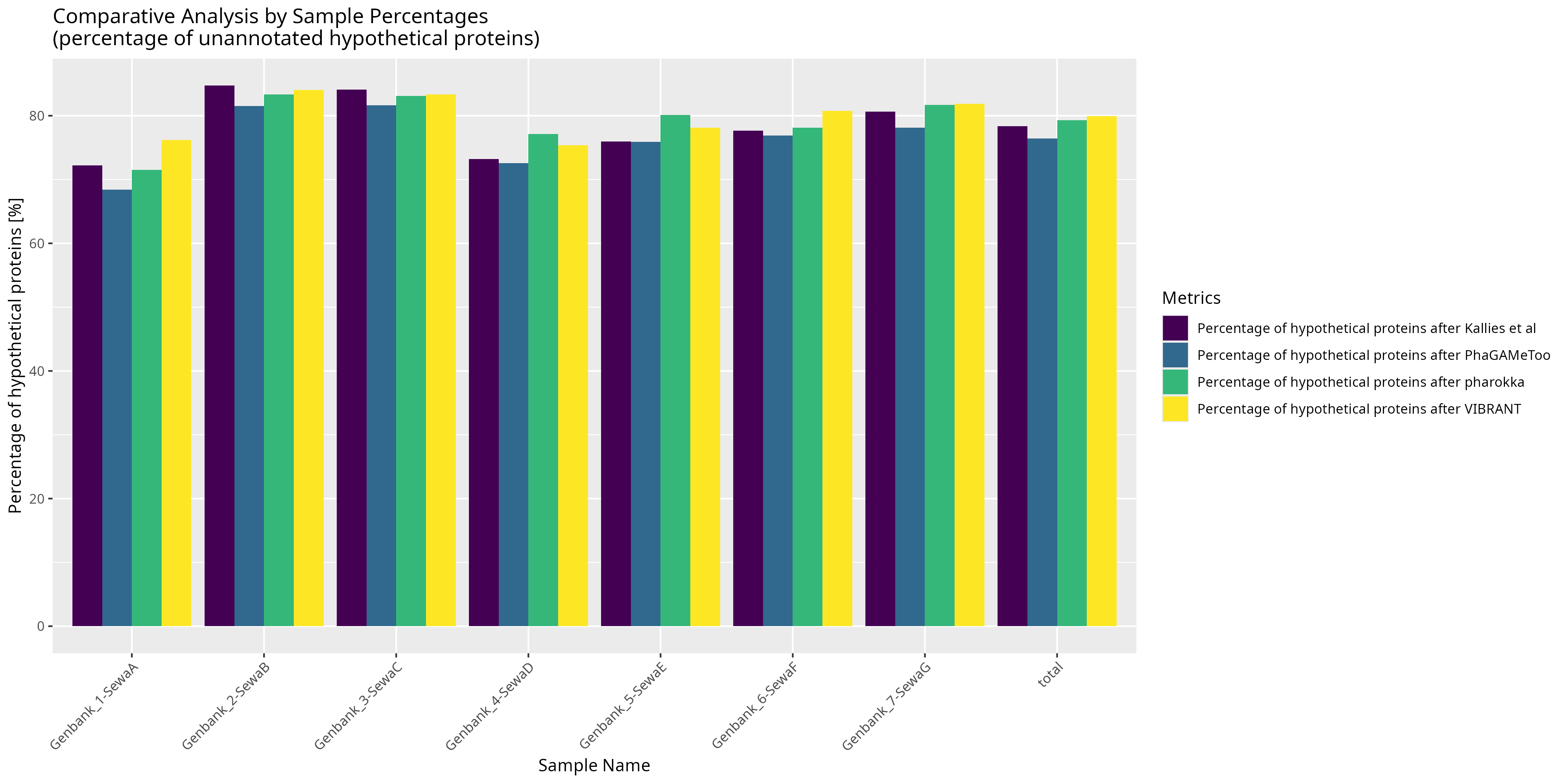
